# AlleleAnalyzer: a tool for personalized and allele-specific sgRNA design

**DOI:** 10.1101/342923

**Authors:** Kathleen C. Keough, Svetlana Lyalina, Michael P. Olvera, Sean Whalen, Bruce R. Conklin, Katherine S. Pollard

**Affiliations:** Pharmaceutical Sciences and Pharmacogenomics Graduate Program at the University of California, San Francisco, California, USA; Gladstone Institutes, San Francisco, California, USA; Bioinformatics Graduate Program at the University of California, San Francisco, California, USA; Departments of Biostatistics, Medicine, Ophthalmology and Pharmacology, University of California, San Francisco, California, USA; Department of Epidemiology & Biostatistics, Institute for Human Genetics, Quantitative Biology Institute, and Institute for Computational Health Sciences, University of California, San Francisco, California, USA; Chan Zuckerberg Biohub, California, USA

**Keywords:** CRISPR, sgRNA design, genomics, genome surgery, genome editing, computational biology

## Abstract

The CRISPR/Cas system is a highly specific genome editing tool capable of distinguishing alleles differing by even a single base pair. However, current tools only design sgRNAs for a reference genome, not taking into account individual variants which may generate, remove, or modify CRISPR/Cas sgRNA sites. This may cause mismatches between designed sgRNAs and the individual genome they are intended to target, leading to decreased experimental performance. Here we describe AlleleAnalyzer, a tool for designing personalized and allele-specific sgRNAs for genome editing. We leverage >2,500 human genomes to identify optimized pairs of sgRNAs that can be used for human therapeutic editing in large populations in the future.

## Background

The CRISPR/Cas genome-editing system is highly specific, with the ability to discriminate between similar genomic sites, even alleles, based on a single nucleotide difference[1]. In order to target a genomic region with the CRISPR system, a single-guide RNA (sgRNA) must be designed that is specific to the region of interest. While current sgRNA design tools incorporate various data relating to predicted efficiency and specificity such as epigenetic marks and chromatin accessibility[2–4], in the vast majority of cases, sgRNAs are designed using reference genomes, such as the hg38 assembly for human or the GRCm38 assembly for mouse. Since sgRNAs are often used on cell lines or organisms with many nucleotide differences from the reference (e.g., on average 0.1% of a human genome[5]). Despite the finding that sgRNAs can sometimes tolerate a single basepair mismatch, these mismatches frequently negatively impact sgRNA efficiency and render imprecise the results of specificity prediction[2, 6, 7]. Furthermore, the use of CRISPR to research areas such as haploinsufficiency, genomic imprinting, and dominant negative diseases require allele-specific sgRNA design. To address these challenges, we developed AlleleAnalyzer, a software tool that designs personalized and allele-specific sgRNAs for individual genomes, identifies pairs of sgRNAs to generate excisions likely to block expression of a gene, and leverages patterns of shared variation from >2,500 human genomes to design sgRNA pairs for that will have the greatest utility in a target population.

## Results and Discussion

Incorporating genetic variation into sgRNA design enables personalized and allele-specific CRISPR experiments. Personalized design involves accounting for variants that disrupt, generate or modify sgRNA sites in a given genome. A genetic variant can impact sgRNA sites by being located in or near a protospacer adjacent motif (PAM site), potentially generating or eliminating sgRNA sites in an individual in a heterozygous or homozygous manner. Rather than being an impediment, these variants can be incorporated into sgRNA design, yielding personalized or allele-specific sgRNAs, depending on variant zygosity (Figure 1a). Because Cas nucleases have different PAM sequences, a variant may impact an sgRNA site for one Cas but not another. We analyzed 11 Cas types (Supplementary Table 1) and ~81 million genome-wide variants annotated by the 1000 Genomes Project[8] (1KGP), finding that most variants impact sgRNA sites for at least one Cas type, even when considering only variants in PAMs, which are putatively more allele-specific[1] (Figure 1b). The likelihood that a variant impacts an sgRNA site differs across Cas nucleases (range: 19-98%), is positively correlated with PAM frequency in the reference genome (Pearson rho=0.9, p=0.04), and is negatively correlated with PAM size (Pearson rho=-0.9, p=0.05). In fact, 3.6% of sgRNAs in the widely used Brunello genome-wide CRISPR screening sgRNA library[9] contain at least one common genetic variant (AF > 5% in the 1KGP cohort), and 2.1% of these sgRNAs contain a variant in the individual human genome of an induced pluripotent stem cell (iPSC) line WTC, commonly used for disease modeling [10] (Figure 1c), impacting ~13% of protein-coding genes in both cases. Failing to account for variants can reduce the efficacy of sgRNAs and also generate unexpected off-target effects. These results emphasize the importance of designing sgRNAs using the personal genome of the patient or cell line where they will be deployed, or at least accounting for both heterozygous and homozygous genetic variants when interpreting results using generic sgRNA libraries.

**Fig. 1.**
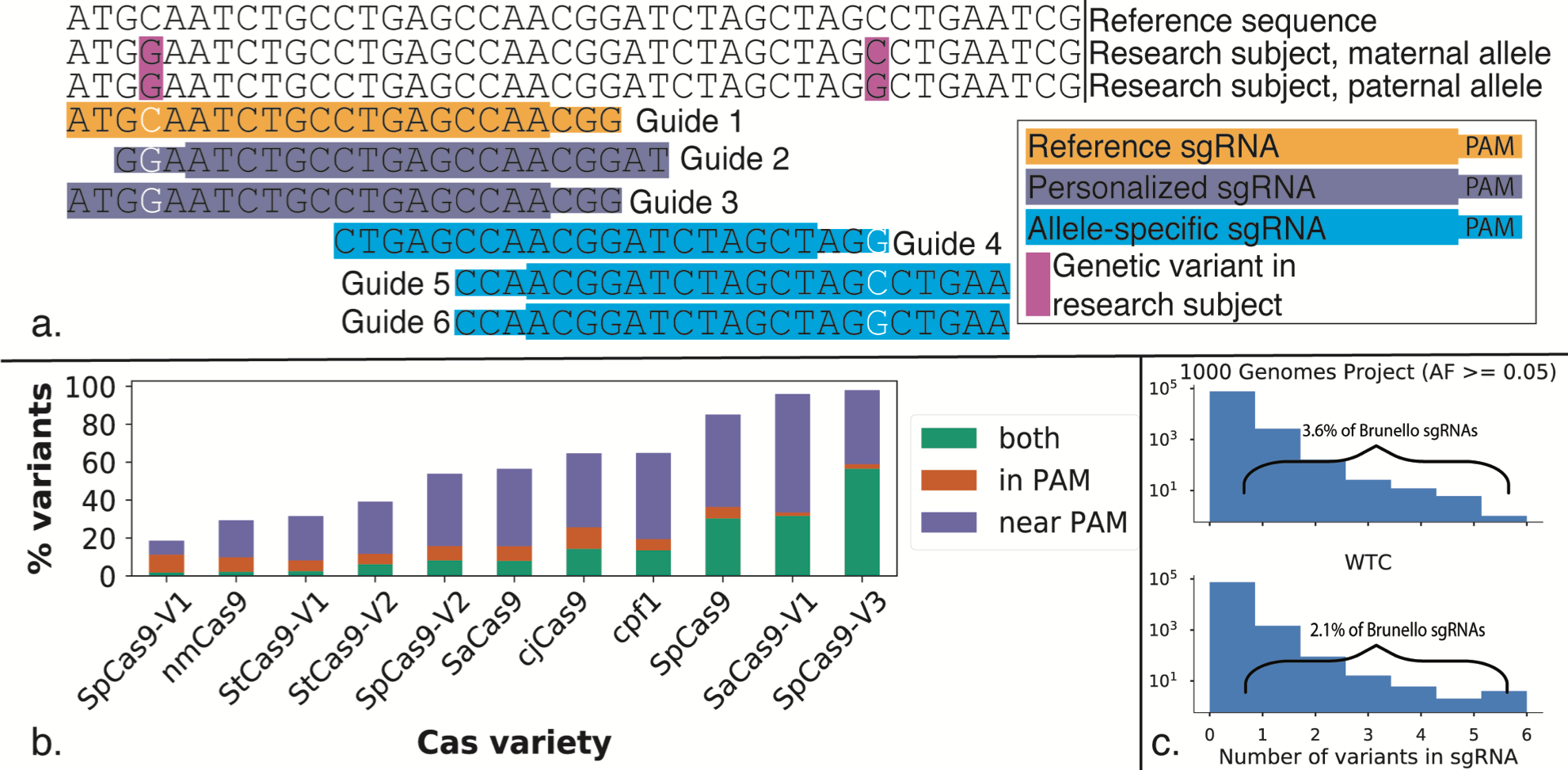
Analysis of allele specific sgRNA sites. In a sample genome, tools designing sgRNAs for the reference genome are imperfect matches due to genetic variants, exemplified by guide 1. This tool designs personalized sgRNAs, as demonstrated by guides 2 and 3, which incorporate homozygous and avoid heterozygous variants. It also designs allele-specific sgRNAs based on incorporation of heterozygous variants, shown by guides 4-6. B) Most variants annotated by the 1000 Genomes Project (1KGP) are in or near a PAM site, demonstrating both a need as well as an opportunity for sgRNA personalization. C) Analysis of common variants, and variants in an individual cell line within the Brunello sgRNA library.

Genetic variants are not just an impediment to sgRNA design; they can be leveraged to establish new therapeutic and research possibilities. Questions that allele-specific editing could help address include haploinsufficiency, imprinting, and allele-specific gene regulation, as well as discovery and correction of heterozygous disease variants. One promising example is genome surgery to treat dominant negative disease by excising only the disease causing copy of a gene, an approach which rescues healthy phenotypes in cell and animal models of dominant negative diseases including Huntington’s disease[11] and retinitis pigmentosa[12, 13]. We assessed this strategy genome-wide by attempting to design a pair of allele-specific sgRNAs for each human protein-coding gene that could generate a genomic excision and eliminate protein production from just one allele. Given a Cas nuclease, an estimated maximum distance between the two sgRNAs on the haplotype to be excised, and allele-specific sgRNA sites, it is possible to classify genes–or other genomic elements, such as enhancers–as putatively targetable or not (Supplementary Figure 1). We use the term putatively targetable when a pair of allele-specific sgRNAs exists but has not yet been tested, because it will not always be possible to cut specifically at a site and coding exon excision will not always stop expression[14]. If we choose a maximum distance of 10 kilobases (kb) between sgRNAs, require the sgRNAs to be within the gene including introns, and consider 11 Cas varieties, the average individual from 1KGP is putatively targetable for allele-specific excision at 77% of protein-coding genes. This rate is evenly distributed across chromosomes but varies by Cas nuclease and gene (Supplementary Figure 2). For genes that are not putatively targetable, additional allele-specific sgRNA sites may be found by leveraging non-coding variants up- and down-stream of the gene, or even in distal enhancers for the gene. Genome-wide, we found that by simply including the 5 kb flanking regions of each gene, we can increase the expected proportion of targetable protein-coding genes per individual from 77% to 85%. We conclude that allele-specific excision is applicable to the vast majority of genes in most human genomes.

**Fig 2:**
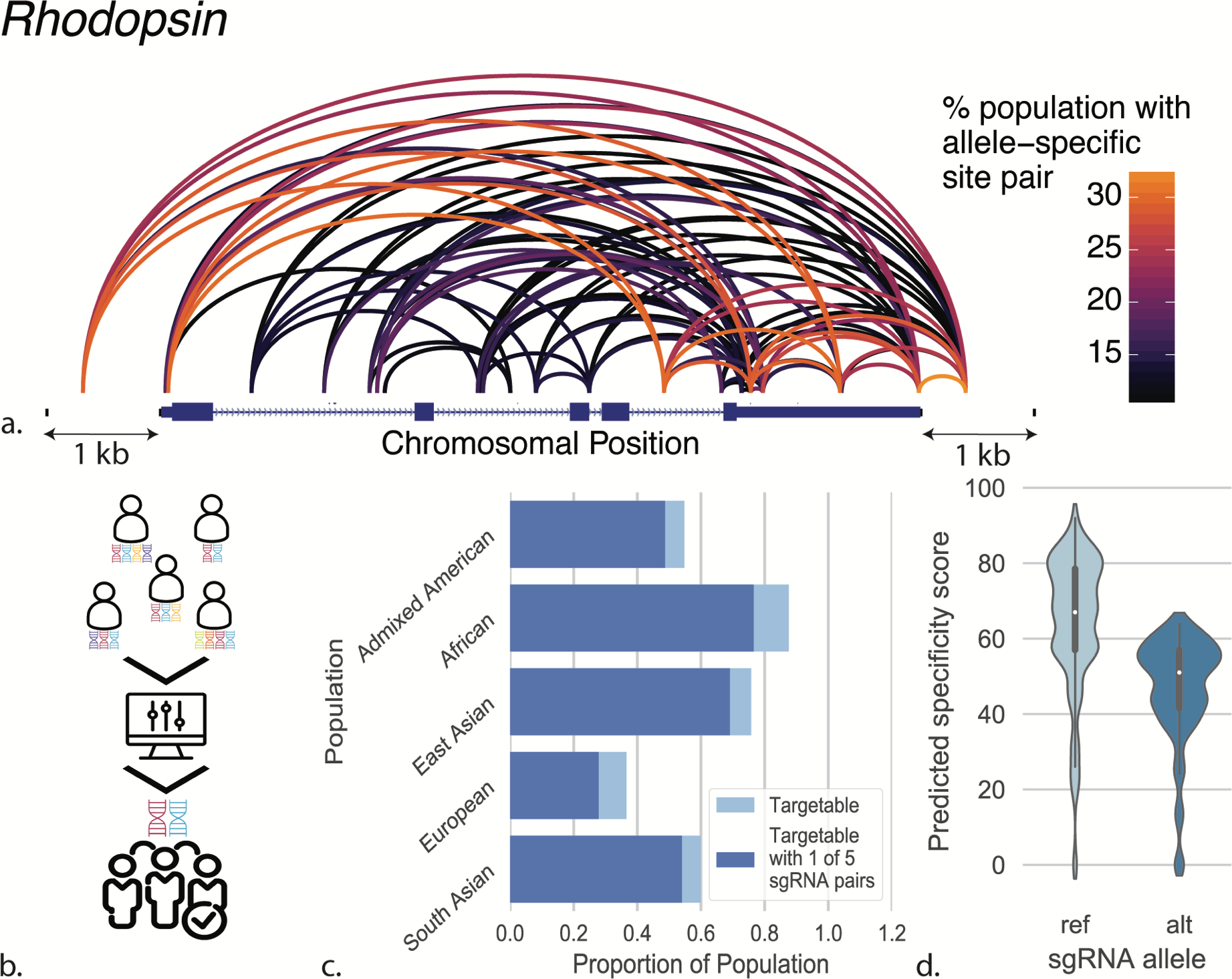
Targeting pairs of allele specific polymorphisms A)82 pairs of allele-specific sgRNA sites for SpCas9 are shared by at least 10% of 1KGP in the gene RHO including 1 kb flanking the gene. B) Development of a set cover optimization algorithm allows targeting of the largest population possible with the fewest allele-specific sgRNA pairs. C) Most targetable individuals have at least one of five SpCas9 sgRNA pairs optimized to be highly shared among the 1KGP cohort in the gene RHO plus the 1 kb regions flanking the gene. D) All possible allele-specific sgRNAs for SpCas9 in RHO plus the flanking 1 kb sequences were designed and scored for predicted specificity using CRISPOR. In the violinplot showing the score distribution for sgRNAs designed against the reference and alternate alleles per each heterozygous variant, the inner boxplot denotes the quartiles of each dataset.

Since some genes in a given individual do not have a pair of allele-specific sgRNAs, we asked if gene silencing with a single allele-specific sgRNA within the coding sequence (single-guide strategy) makes more genes excisable. We compared paired-guide and single-guide strategies for allele-specific gene knockout in the individual human genome of the WTC iPSC line [10] and found that more than twice as many genes are putatively targetable with paired guides (Supplementary Figure 3), because one or both sgRNAs can fall in introns or untranslated regions whereas single sgRNAs are limited to coding regions. Genes that are putatively targetable with a single- and not paired-guide approach tend to have less than two heterozygous variants in the gene, indicating lack of multiple variants as the primary reason a paired-guide strategy fails. These genes likely could be putatively targetable with a paired-guide strategy by incorporating flanking, promoter, or other regulatory regions. We therefore recommend paired-guides for allele-specific gene excision.

Genome editing sgRNAs do not need to be designed one genome at a time. Variants that impact sgRNA sites are often shared among large proportions of the individuals within and sometimes between populations due to haplotype structure. Allele sharing varies by population and locus, as individuals with common ancestry will share haplotypes that harbor specific sets of variants. We therefore developed an algorithm to identify allele-specific sgRNA guide pairs for a given gene that cover the maximum number of individuals in a population; these have the broadest therapeutic potential, similar to designing a drug to treat as many people as possible. Specifically, our method seeks to cover the most people with the fewest sgRNA pairs using their shared heterozygous variants; this is similar to the set cover problem in that the algorithm identifies an optimal combination rather than simply selecting most shared sgRNA pairs, which could disproportionately favor one group over another [15]. Our algorithm generates optimized pairs of sgRNAs that can be used to study or treat genetic diseases in large groups, potentially eliminating the need to develop new sgRNA pairs for each patient or cell line, with practical implications for the development of genome surgery as a field. Our algorithm can also be used to identify sgRNA pair combinations applicable to a custom cohort, enabling researchers to design guides that are maximally shared among multiple cell lines, for example, which would improve experimental efficiency.

As a case study, we investigated the feasibility of excising one allele of exon 1 of *RHO,* which can cause dominant negative macular dystrophy[13]. Considering the gene plus 1 kb of flanking sequence on either side, there are 82 pairs of allele-specific sgRNA sites for SpCas9 that are shared by >10% of all 1KGP individuals, with the number and composition of these pairs varying across 1KGP populations (Figure 2a, Supplementary Figure 4). We sought to identify an optimal combination of five allele-specific sgRNA pairs to target the majority of the 1KGP cohort (Figure 2b). We found that five allele-specific sgRNA pairs could putatively excise one allele of *RHO* while leaving the other allele intact in ~88% of 1KGP individuals with at least two variants, or 57% of the overall 1KGP population (Figure 2c). We also demonstrated how avoiding heterozygous variants and incorporating homozygous variants enables personalized sgRNA design in the *RHO* locus for the WTC genome for many Cas varieties, including SpCas9, SaCas9 and cpf1 (Cas12a) (Supplementary Figure 5, Supplementary Tables 2 and 3). The dominant negative disease gene *RHO* clearly demonstrates the power of using genetic variation in sgRNA design.

We incorporated these methods into AlleleAnalyzer, an open-source software tool (Supplementary Figure 6). This tool designs personalized and allele-specific sgRNAs for unique individuals and cohorts, given their genetic variants, and optimizes sgRNA pairs to cover many individuals based on shared variants. To our knowledge, this is the first computational resource that designs personalized and allele-specific CRISPR sgRNAs, thus expanding and building upon the existing repertoire of sgRNA design tools (Supplementary Table 4). We integrated the specificity scoring capabilities of CRISPOR[4] to enable users to stratify guides by that metric as desired (Figure 2d). The AlleleAnalyzer toolkit and tutorials are available along with the database of annotated 1KGP variants (Supplementary Table 5) at https://github.com/keoughkath/AlleleAnalyzer.

## Conclusions

The genetic variation aware sgRNA design tool AlleleAnalyzer is an important step towards effective deployment of CRISPR-based technologies in diverse genomes, including but not limited to research and therapeutic development for once incurable dominant negative diseases.

## Methods

### PAM occurrence in the human reference genome

#### PAM frequency

The AlleleAnalyzer tool includes a script enabling scanning of a reference genome fasta file for existing PAM sites. We used this to identify PAM sites for 11 Cas types (Supplementary Table 1) in the reference human genomes hg19 and hg38.

### PAM size

PAM sizes were equated as the sum of non-N (A, C, G or T) bases in a PAM site. Thus “NGG” for SpCas9 would have size 2, and “NNGRRT” for SaCas9 would have size 4.

### AlleleAnalyzer analysis of the 1000 Genomes cohort

#### Annotation of variants

Genetic variants were determined to generate or destroy an allele-specific sgRNA site if they were proximal to or in a PAM site (Figure 1a). Sufficient proximity to a PAM site was defined for this study as 20 base pairs based on the common length of sgRNA recognition sequences. For all Cas varieties this was the 20 base pairs 5’ of the PAM, except for cpf1 (Cas12a) for which it was 3’ of the PAM. The sgRNA design tools that are part of AlleleAnalyzer allow different user-defined sgRNA lengths and addition of Cas enzymes and PAMs. There is evidence to suggest that genetic variants that generate or destroy a PAM are more likely to lead to allele-specific Cas activity compared to those in the seed sequence^1^; AlleleAnalyzer thus provides options to differentiate between CRISPR sites in a PAM site versus the sgRNA recognition sequence. All variants genome-wide were annotated for the 1KGP cohort for reference genomes hg19 nd hg38 and are available for querying; an example subset of these data for the first 100 variants annotated by 1KGP on chromosome 1 in reference genome hg19 is available in Supplementary Table 5.

#### Generation of gene set

The gene set analyzed was compiled using the canonical transcripts for RefSeq gene annotations for human reference genome hg19 and hg38 downloaded using the UCSC table browser[16], and filtered for genes with at least one coding exon. When non-protein-coding genes were excluded, 15,199 genes were evaluated for hg19, and 16,143 for hg38. Values reported in the text are for hg19 unless stated otherwise, but analyses were conducted for both reference genomes with similar results.

#### Allele-specific putative gene targetability genome-wide

Putative allele-specific targetability of a gene is defined here as whether a gene contains a pair of allele-specific sgRNA sites for at least one of the 11 Cas enzymes evaluated that are less than 10 kb apart on the same haplotype in an individual that will disrupt a coding exon (Supplementary Figure 1). This metric was calculated for each protein-coding gene for all 2,504 1KGP individuals.

#### Set cover analysis

In order to determine optimal pairs of sgRNAs to cover large groups of people in a particular gene, we applied set cover optimization which we implemented using the Python package PuLP[17]. The aim was to maximize the number of individuals from the 1KGP for whom a user-supplied maximum number of sgRNA pairs would putatively target a given gene. This script can also be used to determine a minimum percentage of people to be covered by a set of sgRNA pairs.

### WTC sequencing

The genome for the iPSC line WTC[10] was sequenced by the Allen Cell Science Institute. Analysis and variant calls in the reference genome hg19 were done according to GATK version 3.7 best practices[18] and phased using Beagle version 4.1 with default settings[19].

### WTC targetability analysis

Variant annotation procedures were the same as in the 1KGP analysis. The same genes lists used in the 1KGP analysis were analyzed in WTC, except when specified in the text, for the cases of 1 kb flanking the gene *RHO*, or when analyzing targetability for all genes + 5 kb flanking vs. genic region only.

### Packages used

#### Python

Docopt was used for handling of command-line arguments. Pandas[20] version 0.21.0 and NumPy[21] version 1.13.3 and elements of the standard Python distribution sys, os, and regex were used for multiple aspects of data analysis. PuLP[17] version 1.6.8 was used for set cover analysis. PyTables[22] was used for data management. Biopython[23] and pyfaidx[24] were used for Fasta processing. Scripts from CRISPOR[4] were integrated into AlleleAnalyzer to facilitate specificity scoring of sgRNAs.

#### R

Packages used to generate arcplots included viridis version 0.5.1, viridisLite version 0.3.0, igraph version 1.1.2, ggraph version 1.0.0, ggplot2 version 2.2.1, reshape2 version 1.4.3, dplyr version 0.7.4, tidyr version 0.7.2, and readr version 1.1.1.

#### Bioinformatics

Bcftools versions 1.5 and 1.6 were used to manipulate VCF and BCF files.

### Scripts

Scripts were written in Python version 3.6.1, R version 3.3.2 and Bash version 3.2.57.

### Data Availability

1KGP phase 3 data were downloaded from the 1KGP website (http://www.internationalgenome.org/). The reference hg19 and hg38 genome data were downloaded from the UCSC genome browser. The 1KGP analysis dataset has been made available for public access online at (http://lighthouse.ucsf.edu/public_files_no_password/excisionFinderData_public/1kgp_dat/).

### Code Availability

All data processing and analysis scripts as well as the sgRNA design tool are located at github.com/keoughkath/AlleleAnalyzer.

## List of Abbreviations

sgRNA: single-guide RNA
PAM site: protospacer adjacent motif site
1KGP: 1000 Genomes Project
kb: kilobases (1000 genomic basepairs)
iPSC: induced pluripotent stem cell

## Declarations

**Ethics approval and consent to participate**

Not applicable.

### Consent for publication

Not applicable.

### Availability of Data and Material

All data processing and analysis scripts as well as the sgRNA design tool are located at github.com/keoughkath/AlleleAnalyzer. 1KGP phase 3 data were downloaded from the 1KGP website (http://www.internationalgenome.org/). The reference hg19 and hg38 genome data were downloaded from the UCSC genome browser. The 1KGP analysis dataset has been made available for public access online at (http://lighthouse.ucsf.edu/public_files_no_password/excisionFinderData_public/1kgp_dat/). WTC whole-genome sequencing data is made available by the Allen Institute at (https://www.allencell.org/genomics.html).

### Authors’ Contributions

K.C.K, S.W., B.R.C., and K.S.P. conceived the project. K.C.K, S.W., S.L., B.R.C., and K.S.P. designed the experiments, K.C.K, S.L., M.P.O., B.R.C., and K.S.P. analyzed data. K.C.K, B.R.C., and K.S.P. wrote the paper with editing from all other authors.

## Acknowledgements

We are grateful for valuable scientific input from Anthony Moore, and members of the Pollard and Conklin labs. We thank Anders Riutta from the Gladstone Bioinformatics Core for his Github troubleshooting expertise. We are grateful to Maximilian Haeussler for his assistance and feedback with the integration of CRISPOR.

---

**Supplementary Figure 1.**
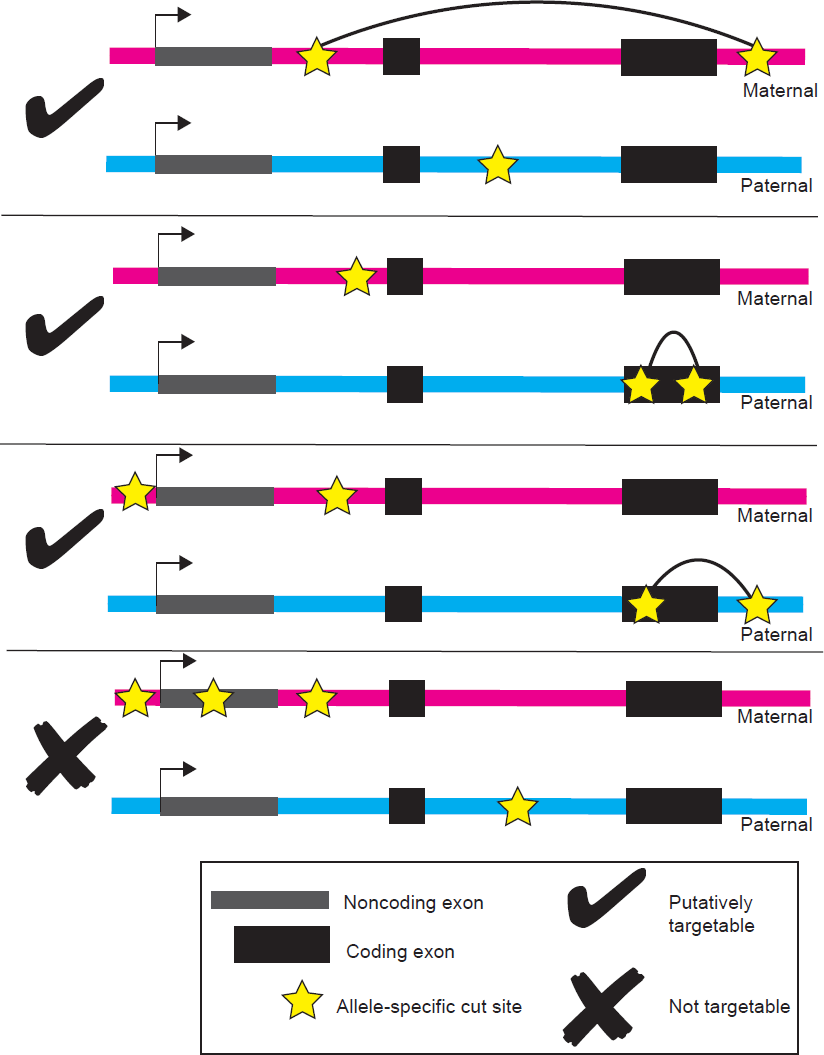
A pair of allele-specific sgRNA sites is defined at putatively targetable if their predicted excision will disrupt at least one protein-coding exon.

**Supplementary Figure 2.**
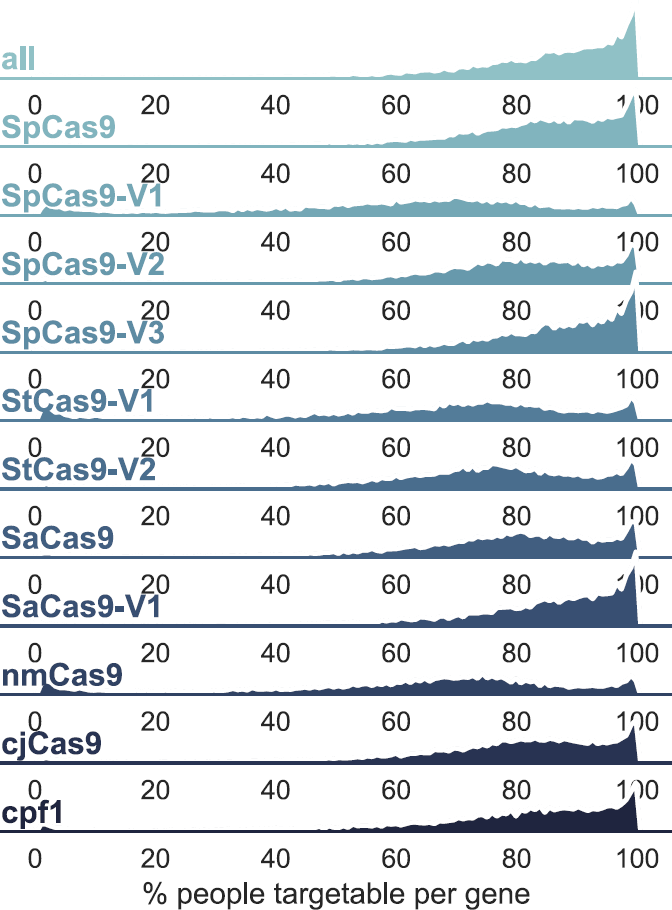
This faceted density plot shows the percentage of putatively targetable 1KGP individuals (2,504 total individuals) per protein-coding gene for 11 types of Cas nuclease.

**Supplementary Figure 3.**
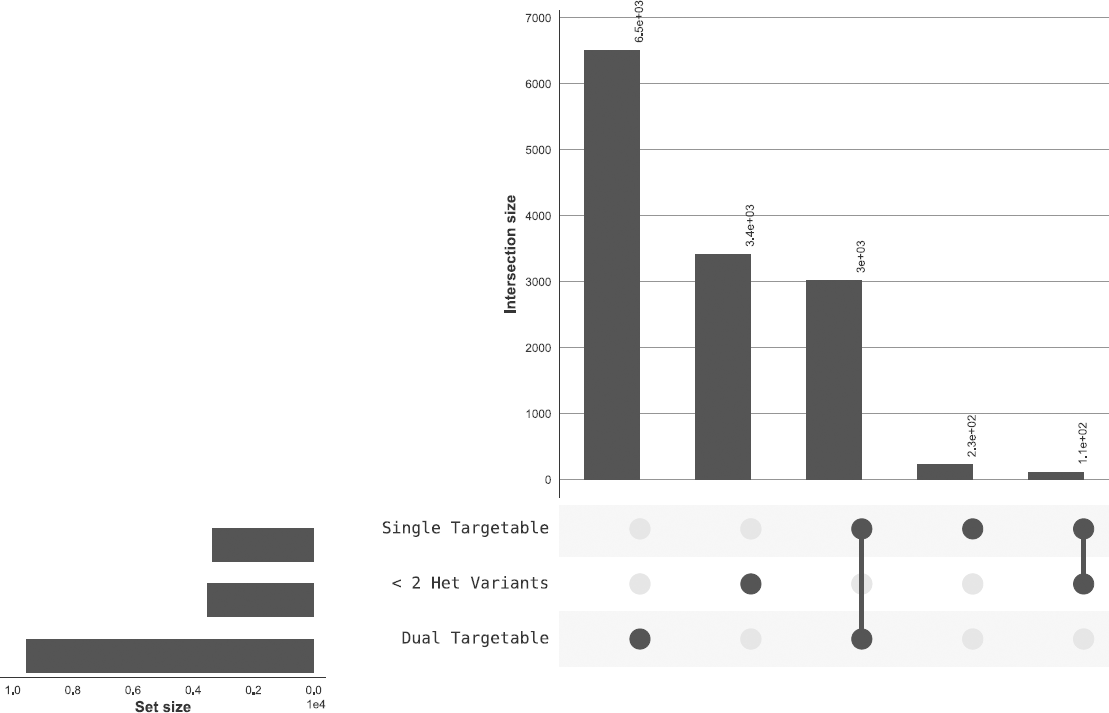
Many more genes are targetable in the genome of WTC with a paired (dual)- as opposed to single-guide strategy. The number of variants in a gene is influential in determining targetability. Many genes that are not dual- or single-guide targetable have very few variants, and the genes that are only targetable with a single-guide approach compared to a dual-guide approach also tend to have fewer variants. All 11 Cas varieties are considered in this analysis.

**Supplementary Figure 4.**
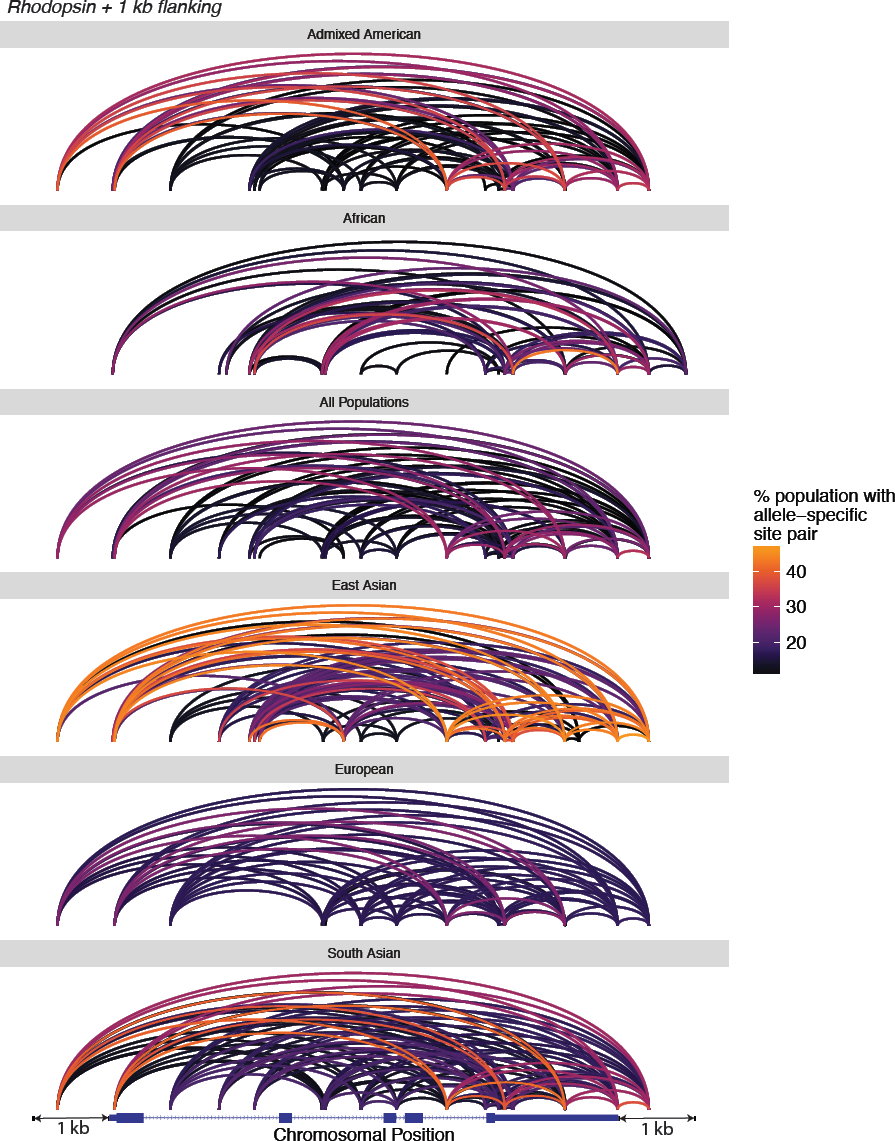
Shared pairs of sgRNAs per locus vary by population. We show allele-specific sgRNA site pairs shared by at least 10% of each population for SpCas9 in the gene *RHO* plus the 1 kb flanking regions in the five super-populations in the 1KGP as well as the overall 1KGP cohort.

**Supplementary Figure 5.**
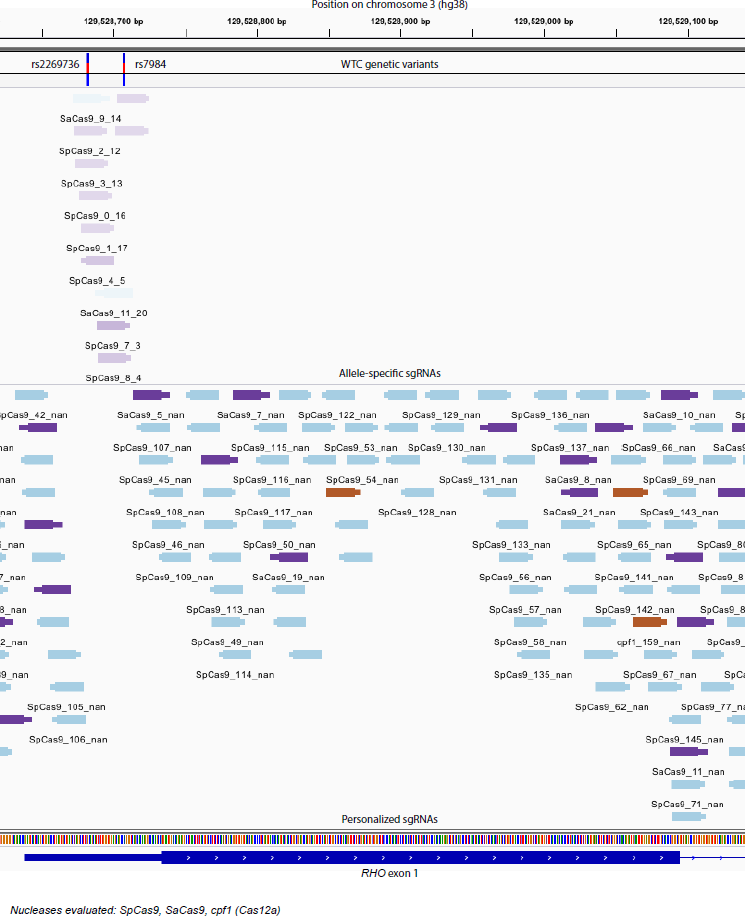
Integrative Genomics Viewer track view of allele-specific (lavender, upper track) and personalized (multi-colored, middle track) sgRNAs for SpCas9, SaCas9 and cpf1 (Cas12a) in the gene *RHO* plus 1 kb flanking in WTC. Allele-specific guides are shaded according to position of the variant in the guide, with variants closer to the PAM being darker based on their putative greater specificity. The track labeled “WTC genetic variants” (top) denotes genetic variants in WTC in this locus, of which there are only heterozygous variants. The bottom track shows the RefSeq annotation for the first exon of this gene in hg38.

**Supplementary Figure 6.**
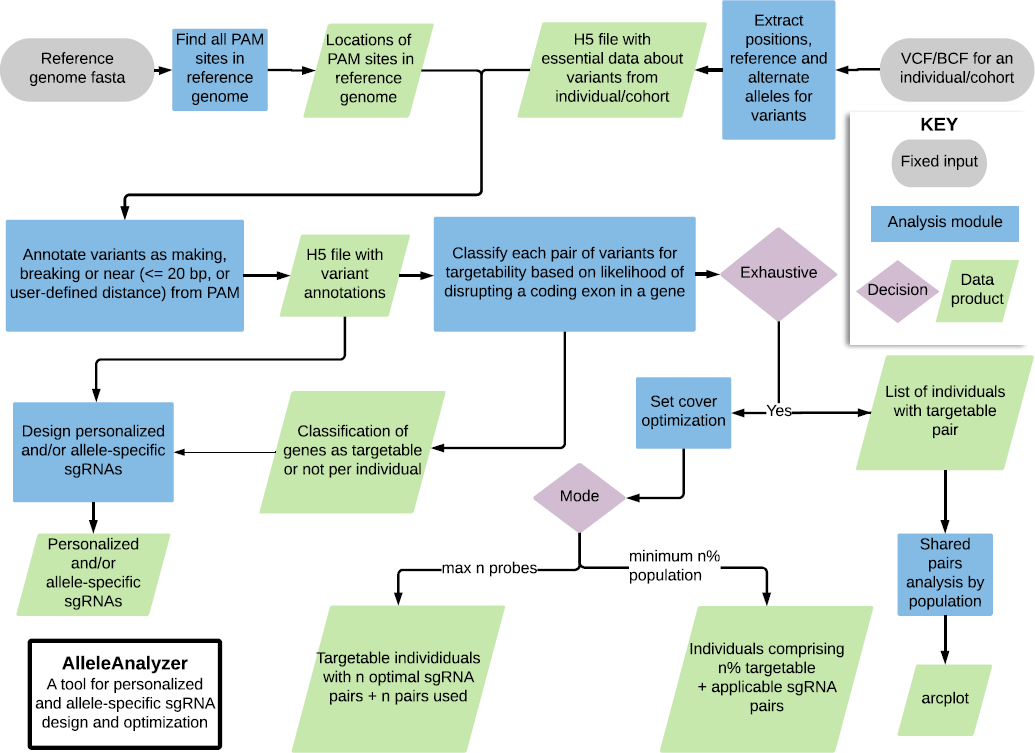
Flowchart for the AlleleAnalyzer software tool.

**Supplementary Table 1.** 11 types of Cas enzyme were evaluated, each of which has a distinct PAM site.

| Common name(s) | Abbreviation | PAM | Properties |
| --- | --- | --- | --- |
| SpCas9 | SpCas9 | NGG | *Streptococcus pyogenes* (Sp) Cas9., most widely used version with dozens of variants using same PAM, e.g. eSpCas9, SpCas9-HF1, eSpCas9 1.1 and more (Jinek et al. 2012) |
| SpCas9 VRER Variant | SpCas9-V1 | NGCG | Version of SpCas9 with alternative targeting range (Kleinstiver et al. 2015) |
| SpCas9 EQR Variant | SpCas9-V2 | NGAG | Version of SpCas9 with alternative targeting range (Kleinstiver et al. 2015) |
| SpCas9 VQR Variant | SpCas9-V3 | NGAN or NGNG | Version of SpCas9 with wider targeting range (Kleinstiver et al. 2015) |
| SaCas9 | SaCas9 | NNGRRT | *Staphylococcus aureus* (Sa) Cas9. Small relative to SpCas9, (Horvath et al. 2008, Jiang et al. 2013) |
| SaCas9 KKH Variant | SaCas9-V1 | NNNRT | Version of SaCas9 with 2 to 4-fold increased targeting range relative of SaCas9 (Kleinstiver et al. 2015) |
| nmCas9 | nmCas9 | NNNNGATT | *Neisseria meningitidis* (Nm) Cas9, with different PAM site (Hou et al. 2013) |
| cpf1 | cpf1 | TTTN | Multiple variations, notably opposite orientation system and sticky-end cut rather than blunt. Multiple species exist, including from *Acidaminococcus* and *Lachnospiraceae*. (Zetsche et al. 2015) |
| StCas9 1 | StCas9-V1 | NNAGAA | *Streptococcus thermophilus* (St) Cas9. Smaller relative of SpCas9. Increased specificity. (Kleinstiver et al. 2015, Muller et al. 2016) |
| StCas9 2 | StCas9-V2 | NGGNG | *Streptococcus thermophilus* (St) Cas9. Smaller relative of SpCas9. Increased specificity. (Muller et al. 2016) |
| cjCas9 | cjCas9 | NNNNACA | *Campylobacter jejuni* Cas9. Smallest Cas9 ortholog to date, easy to package (Kim et al. 2017) |

**Supplementary Table 2** All possible allele-specific sgRNAs for SpCas9, SaCas9 and cpf1 (Cas12a) in the region surrounding the first exon of *RHO* WTC (Supplementary Figure 6).

**Supplementary Table 3.** All possible personalized sgRNAs for SpCas9, SaCas9 and cpf1 (Cas12a) in the region surrounding the first exon of *RHO* WTC (Supplementary Figure 6). WTC has no homozygous variants in this region, thus allele frequency and variant-related columns are blank. However, the sgRNAs are designed to avoid the 9 heterozygous variants that WTC has in this region.

|  | Personalized sgRNA design | Allele-specific sgRNA design | Supports any genome | Paired sgRNA design | Optimizes pairs of sgRNAs to maximize coverage in a cohort | Single or multi-locus design | Multiple Cas varieties | Ability to add novel Cas enzymes | Off-target scoring |
| --- | --- | --- | --- | --- | --- | --- | --- | --- | --- |
| AlleleAnalyzer | yes | yes | yes | yes | yes | yes | yes | yes | yes |
| CRISPOR (Haeussler et al. 2016) | no | no | yes | no | no | yes | yes | no | yes |
| GuideScan (Perez et al. 2017) | no | no | no | yes | no | yes | yes | no | yes |
| E-CRISP (Heigwer et al. 2014) | no | no | no | yes | no | yes | no | no | yes |
| MIT design tool (Hsu et al. 2013) | no | no | no | no | no | yes | no | no | yes |
| CRISPRscan (Moreno-Mateos et al. 2015) | no | no | no | no | no | no | yes | no | yes |
| FlashFry (McKenna & Shendure, 2018) | no | no | yes | no | no | yes | yes | no | yes |

**Supplementary Table 4** Comparison of AlleleAnalyzer features with other commonly used CRISPR sgRNA design tools.

**Supplementary Table 5** An example subset of variant annotations for the first 100 variants on chromosome 1 from 1KGP.

